# Substrate binding allosterically relieves autoinhibition of the TRIB1 pseudokinase

**DOI:** 10.1101/313767

**Authors:** Sam A. Jamieson, Zheng Ruan, Abigail E. Burgess, Jack R. Curry, Hamish D. McMillan, Jodi L. Brewster, Anita K. Dunbier, Alison D. Axtman, Natarajan Kannan, Peter D. Mace

## Abstract

**One Sentence Summary:** Substrate binding to Tribbles-homolog 1 (TRIB1) pseudokinase induces allosteric changes that allow formation of a complex with the COP1 ubiquitin ligase.

**Abstract:** The Tribbles family of pseudokinases recruit substrates to the COP1 ubiquitin ligase for ubiquitination. CCAAT-enhancer binding protein (C/EBP) family transcription factors are crucial Tribbles substrates in adipocyte and myeloid development. Here we show that the TRIB1 pseudokinase can recruit various C/EBP family members, with binding of C/EBPβ attenuated by phosphorylation. To explain the mechanism of substrate recruitment, we solved the crystal structure of TRIB1 in complex with C/EBPα. TRIB1 undergoes a significant conformational change relative to its substrate-free structure, to bind C/EBPα in a pseudo-substrate-like manner. Crucially, substrate binding triggers allosteric changes that link substrate recruitment to COP1 binding, which is consistent with molecular dynamics and biochemical studies. These findings offer a view of pseudokinase regulation with striking parallels to *bona fide* kinase regulation— via the activation loop and αC-helix—and raise the possibility of small molecules targeting either the activation loop-in, or loop-out, conformations of Tribbles pseudokinases.

## Introduction

Protein kinases are a pervasive class of signalling protein that transduce all manner of biological signals. Kinase catalytic activity is controlled by a range of mechanisms, which converge on alignment of the catalytic and regulatory spines (1). Amongst the ~500 protein human protein kinase domains, approximately 10% are regarded as pseudokinases because they lack key catalytic features. Sequence variations that can turn a kinase into a pseudokinase include mutation of catalytic resides, substitutions that block ATP binding, or disruption of regulatory spine residues (2). Like their catalytically active counterparts, pseudokinases play roles in many different signalling pathways, but do so without phosphorylating substrates. These functions can be broadly categorised as allosteric regulation of active kinases, acting as scaffolds for protein– protein interactions, or acting as signalling switches (2, 3).

The Tribbles family of proteins occupy a dedicated branch of the kinome comprised of four members—TRIB1, TRIB2, TRIB3, and the more distantly related STK40 (also known as SgK495) (4). The family derives its name from the Drosophila Tribbles protein (5, 6), and share a common domain architecture: they have a variable N-terminal extension; a pseudokinase domain; and a C-terminal extension that binds to the COP1 ubiquitin E3 ligase (Fig. 1A). The pseudokinase and COP1-binding motif are key for the function of Tribbles proteins—by binding to substrates through their pseudokinase domain and to COP1 through their C-terminus they act as substrate adaptors for ubiquitination by COP1.

**Fig. 1.**
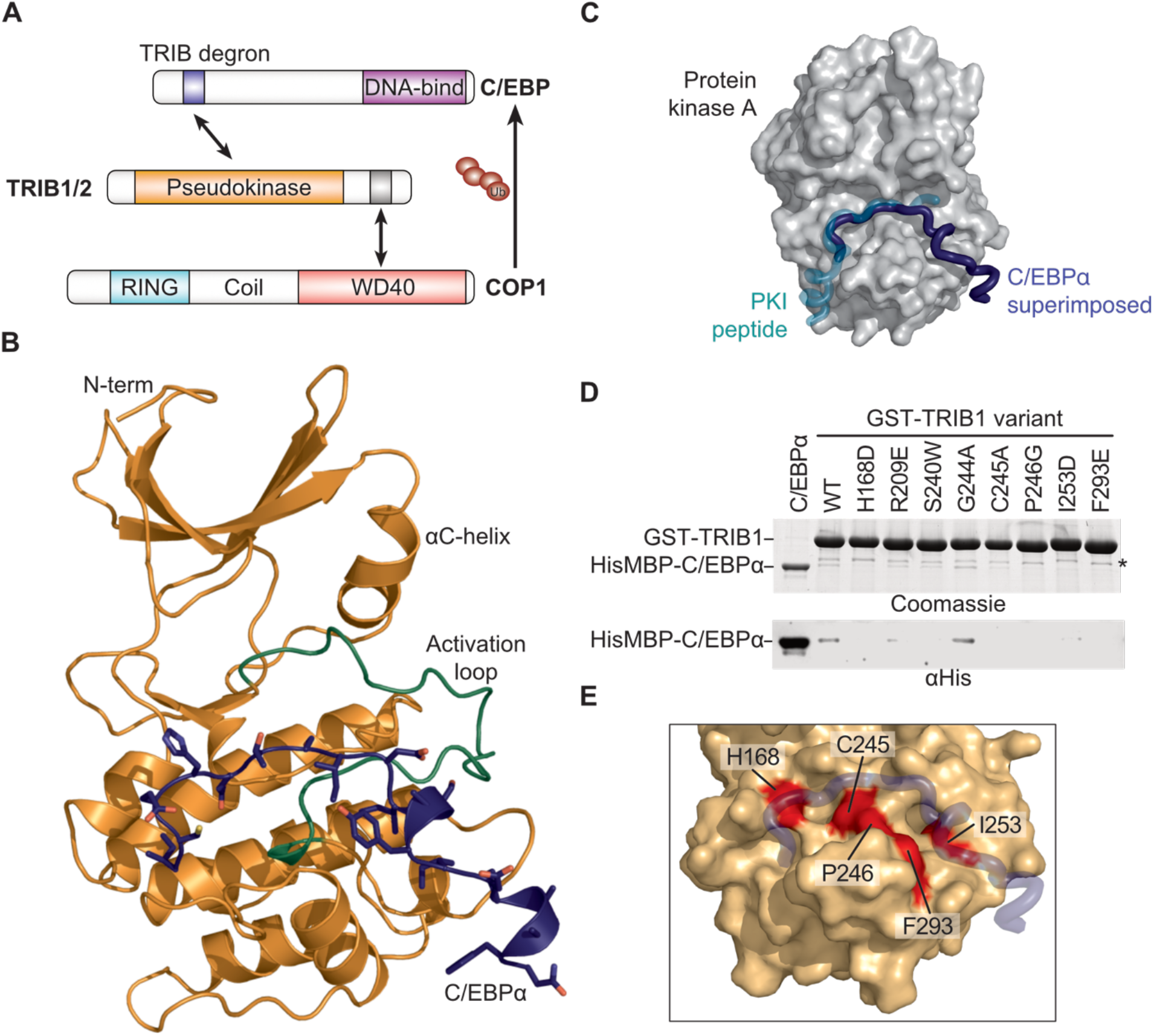
The TRIB1-C/EBPα degron complex. **(A)** Schematic illustrating degradation of C/EBPα by TRIB1-COP1. **(B)** Crystal structure of the TRIB1-C/EBPα degron complex, with TRIB1 shown predominantly in orange with a green activation loop and C/EBPα in blue. **(C)** Comparison of the C/EBPα binding mode (blue) with that of the prototypic substrate-like protein kinase inhibitor (PKI; turquoise) in complex with protein kinase A (grey surface; PDB 1ATP) **(D)** GST-pulldown of His_6_MBP-C/EBPα(53–75) by WT GST-TRIB1(84–372) and indicated mutants, separated by SDS-PAGE and visualised by Coomassie blue staining (upper) or Anti-His_6_ immunoblotting (lower). A non-specific band that co-purifies with GST-TRIB1 is indicated with an asterisk. **(E)** Locations of TRIB1 mutations that disrupt C/EBPα binding, with TRIB1 shown as an orange surface and C/EBPα in blue.

Human Tribbles proteins are implicated in wide-ranging signalling processes. TRIB3 is known to affect metabolism by binding to the metabolic regulatory protein kinase AKT, and recruiting acetyl-CoA carboxylase for degradation (7, 8). In contrast to inducing degradation, TRIB3 can stabilise the oncogenic PML-RARα fusion protein preventing its sumoylation and ubiquitination (9). In addition, TRIB1–3 have all been reported to control MAP kinase signalling (10–13).

Perhaps the most well-established role of Tribbles proteins is regulating CCAAT enhancer binding protein (C/EBP) family transcription factors (14–17). The C/EBP family is comprised of 6 members in humans (α, β, γ, δ, ε, and ζ). Further diversity arises because most family members can be translated as splicing isoforms, and family members can form either homo- or hetero-dimers (18). C/EBP transcription factors recognise a characteristic CCAAT DNA sequence to regulate proliferation, differentiation and metabolism, particularly of hepatocytes, adipocytes and hematopoietic cells (15, 18–20). Amongst the Tribbles, TRIB3 appears to not influence C/EBPs, whereas TRIB1, TRIB2 and STK40 are all capable of triggering degradation of selected C/EBPs by recruiting them to COP1 for ubiquitination (14, 21). Through their ability to control C/EBP and myeloid development, overexpression of TRIB1 and TRIB2 can cause development of acute myeloid leukaemia with high penetrance (14, 22), and TRIB1 deficiency results in a loss of tissue-resident M2-like macrophages (15). GWAS studies have also linked TRIB1 to levels of metabolic enzymes and plasma lipids (23, 24), which has subsequently been linked to post-translational control of C/EBP-regulated transcription (25).

Crystal structures of the TRIB1 pseudokinase domain have revealed a deprecated N-terminal lobe and ATP-binding site, which is consistent with the inability of TRIB1 to bind ATP (17). Importantly, the C-terminal COP1-binding motif of TRIB1 can bind to the pseudokinase domain in an autoinhibitory manner (17), which is mutually exclusive with the motif binding to the COP1 WD40 domain (26). Here we use a combination of crystallography, molecular dynamics and biochemistry to investigate the mechanism that allows release of TRIB1 autoinhibition. The structure of TRIB1 bound to its prototypical substrate, C/EBPα, shows that TRIB1 undergoes a marked conformational change upon substrate binding. Such conformational changes have important implications for assembly of an active complex with COP1, and potentially for susceptibility of Tribbles pseudokinases to small molecule inhibitors.

## Results

### Structure of the TRIB1-C/EBPα complex

To understand the basis for recruitment of C/EBP family transcription factors we sought to solve the structure of the TRIB-C/EBPα complex. Crystals of the complex were successfully grown by fusing the TRIB1 recognition degron of C/EBPα to the C-terminus of TRIB1 (fig. S1). The structure was solved to a resolution of 2.8 Å (Table 1). There are two complexes in the asymmetric unit, which have the C/EBPα peptide defined for residues 55–74 and 57–67, respectively. The former complex is displayed in subsequent figures. Both complexes and electron density maps are included in fig. S2. Remarkably, the activation loop of TRIB1 in both complexes is fully resolved (Fig. 1B), and adopts a position markedly different than the confirmation seen in apo-TRIB1 (17). The activation loop folds towards the αC helix and generates a binding site for the C/EBPα degron. This binding mode is effectively pseudo-substrate-like, as demonstrated by comparison with the prototypical substrate-like protein-kinase inhibitor peptide in complex Protein Kinase A (Fig. 1C). All of the C/EBPα degron residues previously shown to be required for binding make direct contact with TRIB1 (17). Namely Ile55, Glu59, Ser61, Ile62 and Ile 64 are part of an extended peptide that spans TRIB1 and leads to a helical turn containing Tyr67 and Ile68, which lie in a cleft alongside the αG-helix of TRIB1.

**Table 1.** Crystallographic data summary

|  |  |
| --- | --- |
| | TRIB1–cEBPα |
| Beamline | AS-MX2 |
| Wavelength (Å) | 0.9537 |
| Resolution (outer shell) (Å) | 48.7–2.8 (2.95–2.8) |
| Space group | P6<sub>1</sub>22 |
| Unit cell parameters | a = 98.8 Å<br>b = 98.8 Å<br>c = 332.7 Å<br>$\alpha$ = 90°<br>$\beta$ = 90°<br>$\gamma$ = 120° |
| R<sub>merge</sub> (outer shell) | 0.077 (1.992) |
| R<sub>pim</sub> (outer shell) | 0.035 (0.879) |
| Mean I/σI (outer shell) | 19.8 (1.5) |
| Completeness (outer shell) | 99.8 (99.1) |
| Multiplicity (outer shell) | 10.4 (10.7) |
| Total No. of reflections | 258141 (37340) |
| No. of unique reflections | 24777 (3491) |
| Mean (I) half-set correlation CC(1/2)<br>(outer shell) | 1.000 (0.609) |
| Wilson B factor (Å<sup>2</sup>) | 82.1 |
| Refinement statistics |  |
| R<sub>cryst</sub> | 0.220 |
| R<sub>free</sub> | 0.276 |
| Average B -factor overall (Å<sup>2</sup>) | 113 |
| Ramachandran plot statistics (%) |  |
| Favored regions | 93.1% |
| Allowed regions | 6.5% |
| Outliers | 0.4% |
| PDB entry |  |

To experimentally validate the TRIB1 residues involved in the interface, we generated a panel of mutants in a GST-TRIB1 fusion protein, and tested their ability to bind His_6_MBP-C/EBPα (Fig. 1D,E). Mutants were selected that either directly contact C/EBPα or stabilise the activation loop in a conformation that is competent to form the binding site. In pulldown analysis, critical residues span the C/EBPα binding site—either directly contacting C/EBPα (His168, Phe293), or stabilising the substrate-competent conformation of the activation loop (Cys245, Pro246, and Ile253). The five TRIB1 mutants with abrogated C/EBPα binding are identical in TRIB2, consistent with the ability of TRIB1 and TRIB2 to bind C/EBPα with similar affinities (fig. S3), and to subsequently induce degradation of C/EBPα in cells and drive development of acute myeloid leukaemia in mice (14).

### TRIB1 binds multiple C/EBP family members

Amongst their broad functions, C/EBP transcription factors coordinately regulate myeloid cell and adipocyte differentiation, with different family members predominating at specific phases of differentiation. However, there is some sequence variation in the Tribbles recognition degron of different C/EBP transcription factors (Fig. 2A)—C/EBPβ, δ and ε each contain a sequence similar to the C/EBPα degron, while C/EBPγ and ζ do not appear to have a comparable motif. We first sought to test whether C/EBPβ, δ, and ε could also be bound by TRIB1, and subsequently if sequence variations might alter the efficacy of TRIB1 binding. Employing isothermal titration calorimetry, we observed that TRIB1 bound to C/EBPα, β, δ and ε, peptides fused to maltose-binding protein with dissociation constants ranging from 7.4 to 17 μM (Fig. 2B). Such affinities suggest that C/EBPα, β, δ and ε all contain functional TRIB1 degrons, that bind TRIB1 with similar affinity.

**Fig. 2.**
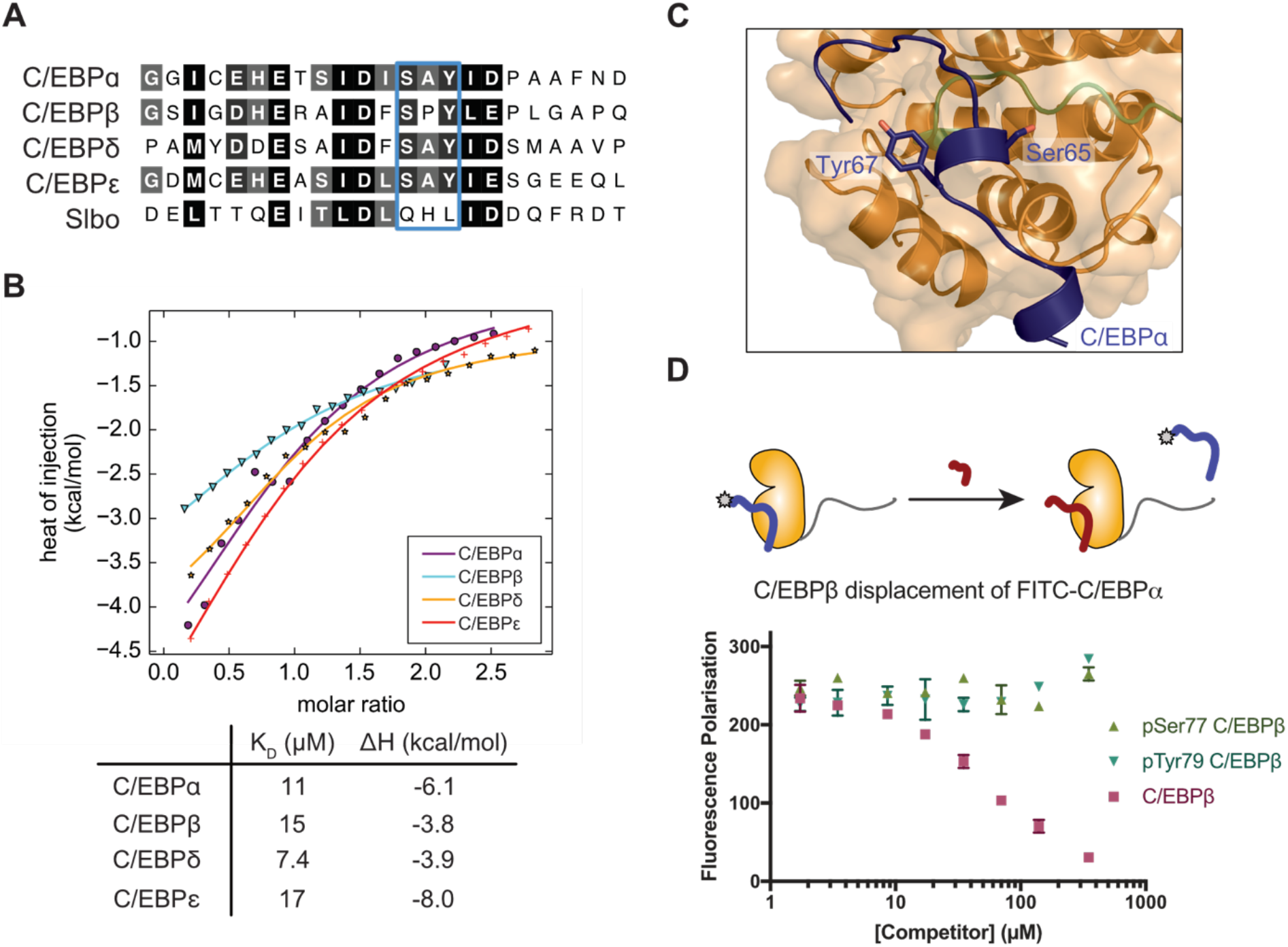
TRIB1 binds various C/EBPs and can be antagonized by phosphorylation. **(A)** Alignment of the TRIB degron from various human C/EBP proteins, and the Drosophila C/EBP ortholog, Slbo. **(B)** Isothermal titration calorimetry of indicated MBP-fused C/EBP degrons injected into TRIB1(84–372). Data points represent mean of duplicate titrations, following buffer titration subtraction. **(C)** Detailed view of Ser65 and Tyr67 within the C/EBPα degron (equivalent to Ser77 and Tyr69 in C/EBPβ) in complex with TRIB1. **(D)** Fluorescence polarisation displacement assay of FITC-C/EBPα degron from TRIB1 by: unmodified, Ser77-phosphorylated, or Tyr79-phosphorylated, C/EBPβ peptide. Data points represent the mean of three technical replicates with error bars indicating SEM.

To investigate which residues within the C/EBP degron contribute most strongly to TRIB1-binding, we performed a 1 μs molecular dynamics simulation based on the TRIB1-C/EBPα structure. Plotting the root mean square fluctuation of C/EBPα peptide reiterates that the stretch from Glu59–Ala72 represents the most stable portion of the peptide and include the main binding determinants (fig. S4). This finding is consistent with several lines of evidence: previous binding studies of mutant C/EBPα peptides revealed a concentration of important binding residues in this region (17); the second complex within the asymmetric unit of the crystal structure was well defined only for residues 57–67; and this segment contains the highest levels of sequence identity between C/EBP family members.

### C/EBPβ phosphorylation antagonises TRIB1 binding

The observation that TRIB1 can bind various C/EBP family members with similar efficacy is interesting in light of previous observation that TRIB1 can affect C/EBPα but not C/EBPβ in certain cases (15), and other scenarios where C/EBPα and C/EBPβ are regulated equally by TRIB1 (27). These observations beg the question of whether further mechanisms control C/EBP degradation depending on cell type or developmental stage. To explore additional layers of regulation, we analysed data from phosphosite and observed that C/EBPβ is phosphorylated at Ser77 and Tyr79 within its Tribbles degron. Tyr79 phosphorylation of C/EBPβ occurs via c-Abl and Arg non-receptor tyrosine kinases, resulting in stabilisation of C/EBPβ (28), while Ser77 is phosphorylated in a Ras-dependent manner (29). To test the effect of C/EBPβ degron phosphorylation, we developed a competition assay based on fluorescence polarisation (Fig. 2D). In this assay, FITC-labelled C/EBPα peptide was competitively displaced by an unmodified C/EBPβ degron peptide. In contrast, Ser77-phosphorylated, or Tyr79-phosphorylated C/EBPβ degron showed no ability to displace FITC-C/EBPα, even up to concentrations of 350 μM. Loss of TRIB1 binding after Tyr79 phosphorylation suggests a direct mechanism to explain post-translational stabilisation of C/EBPβ by c-Abl (28), based on protection from degradation by TRIB1-COP1. Similarly, phosphorylation of Tyr79 by proline-directed Ser/Thr kinases such as CDK2 or MAPKs could represent a further mechanism by which C/EBP transcription levels can be protected from degradation by COP1-Tribbles under specific circumstances (29).

### Conformational changes induced by substrate-binding

Previous structures of TRIB1 have shown that the pseudokinase domain contains an unusual αC-helix that forms a binding site for the C-terminal COP1-binding motif (17). TRIB1 and TRIB2 also contain a unique SLE sequence at the N-terminus of the activation loop—as opposed to DFG that is seen in most active kinases—which can block the putative ATP-binding pocket. The structure of substrate-bound TRIB1 now reveals distinct conformational changes upon substrate binding (Fig. 3). Analysis of these conformational changes delineate a clear allosteric mechanism to link substrate-binding on one side of TRIB1 to release of the COP1-binding motif from its binding site on the αC-helix—in effect release of TRIB1 autoinhibition (Fig. 3 and movie S1).

**Fig. 3.**
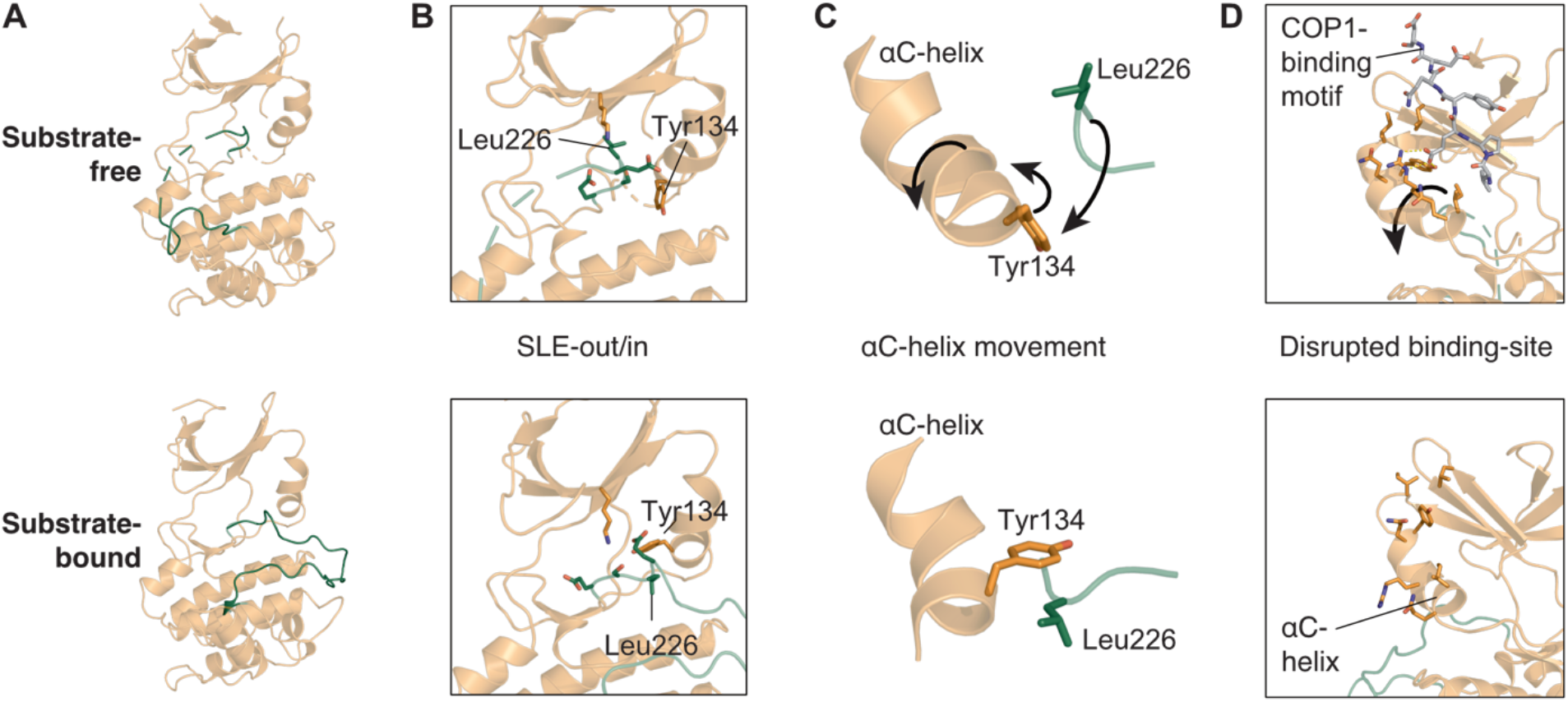
TRIB1 conformational changes upon substrate binding. Comparison of structures of substrate-free (upper panels; PDB 5cem) and C/EBPα-bound TRIB1 (lower panels; this work). **(A)** Overview from the substrate binding side of the molecule. **(B)** The SLE-motif and surrounding residues within the active site, from the same orientation as panel A. **(C)** Simplified view showing the relative organisation of the αC-helix, Tyr134 and Leu226, from the opposite orientation to panels A/B. **(D)** Position of the αC-helix and residues that contact the C-terminal tail of TRIB1 in substrate-free TRIB1; orientation as for panel C. Movement of selected features upon EBPα binding are indicated with arrows in upper panels, and residues are selectively depicted as sticks for clarity.

Upon binding to C/EBPα substrate, the activation loop becomes fully ordered, folding between the N- and C-terminal lobes to form the substrate-binding site (Fig. 3A; Fig. 1). With this rearrangement comes a conformational change in the SLE sequence at the N-terminus of the activation loop (Fig. 3B). Leu226 moves towards the αC helix, packing into a location homologous to the DFG phenylalanine of conventional kinases when they adopt their ‘DFG-in’ conformation (1). In TRIB1, rearrangement of Leu226 is facilitated by movement of Tyr134 within the αC-helix, which subsequently packs atop Leu226 to complete the regulatory spine (Fig. 3C). In this manner, the substrate-bound conformation of the TRIB1 pseudokinase can be thought to adopt a ‘SLE-in’ conformation that is comparable to the ‘DFG-in’ conformation of conventional kinases.

The existence of states that resemble inactive (SLE-out) and active (SLE-in) protein kinases structures is intriguing, but also has striking consequences for the function of TRIB1. In order for co-ordinated rearrangement of Tyr134 and Leu226 upon substrate-binding, the αC-helix of TRIB1 must rotate away from the active site of the kinase. For instance, the α-carbon positions of Tyr134, Ile135 and Gln136 move by 3.2, 4.8 and 4.4 Å, respectively, relative to their positions in the substrate-free structure. Further movement is propagated through the N-terminal portion of the αC-helix, and more subtle rearrangements occur within the β4 and β5 of the N-terminal lobe. Together these changes disrupt the docking site for the C-terminal tail of TRIB1 (Fig. 3D), making it incompatible with sequestration of the C-terminal COP1-binding site and TRIB1 autoinhibition.

### Substrate-binding allosterically regulates TRIB1 autoinhibition

The C-terminal COP1-binding motif of TRIB1 binds to its own pseudokinase domain and the E3 ubiquitin ligase in a mutually exclusive manner (26); (Fig. 4A/B), which necessitates the allosteric changes outlined in the previous section. To test the functional effects of the allosteric changes we adopted an assay previously used to test COP1-binding by TRIB1, and STK40 (26, 30). Namely, TRIB1-COP1 association was inferred by displacement of a FITC-labelled TRIB1 C-terminal-peptide from the WD40 domain of COP1, which can be monitored by fluorescence polarisation. Consistent with Uljon et al. and the autoinhibited crystal structure, we observed that the C-terminal COP1-binding motif is less effective at binding COP1 when it is attached to the pseudokinase (TRIB1(84–372)) than it is as an isolated peptide (residues 349–367; fig. S5). Strikingly, inclusion of the C/EBPα degron peptide with TRIB1(84–372) allows it to more effectively displace the FITC-labelled peptide (Fig. 4C). The displacement curve for TRIB1 with C/EBPα is shifted to approximately 10-fold lower TRIB1 concentrations relative to TRIB1 in the absence of substrate, supporting a model where substrate-binding releases autoinhibition and frees the tail to bind COP1.

**Fig. 4.**
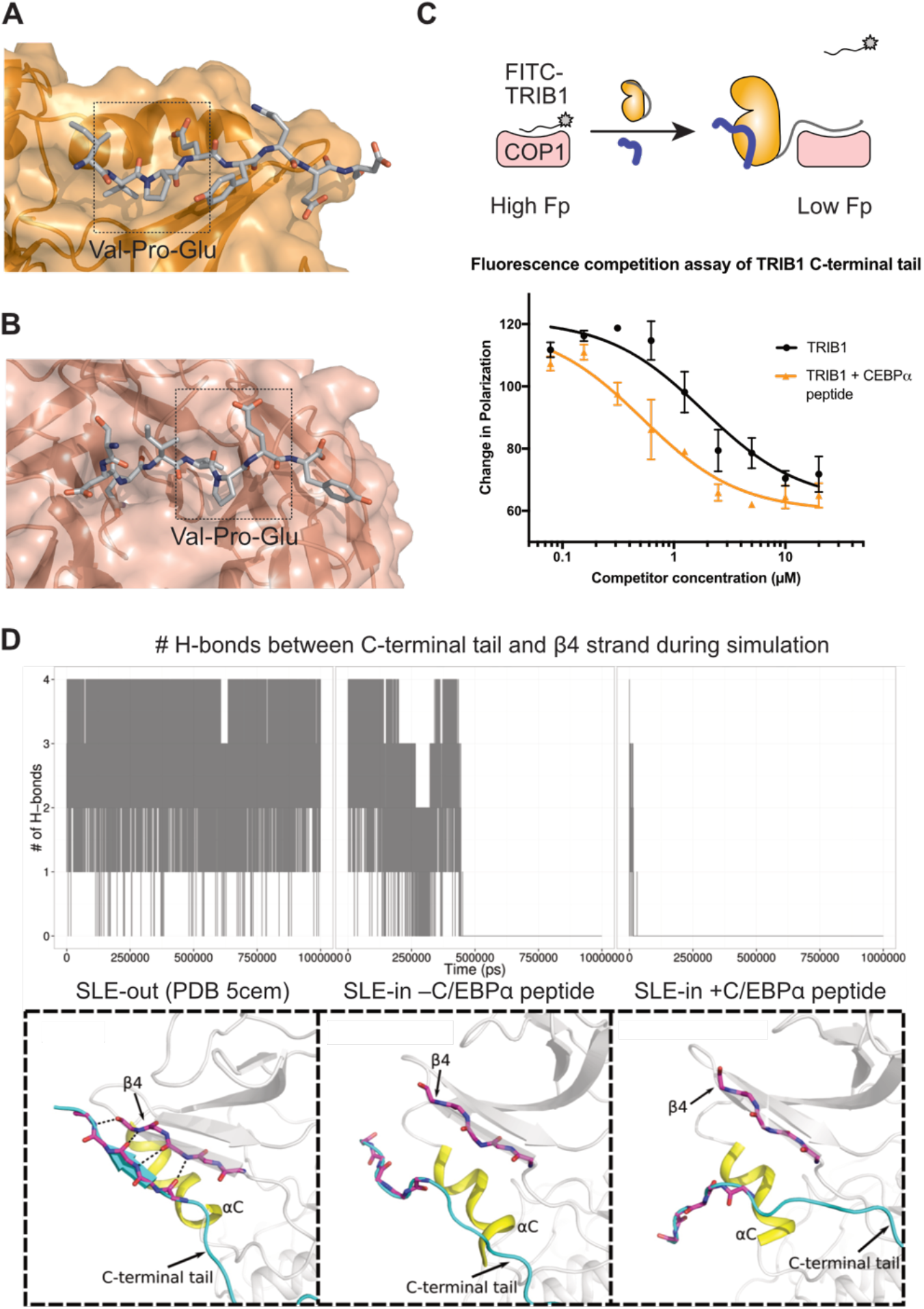
Allosteric release of the COP1-binding motif by substrate-binding. Structure of the C-terminal COP1 binding motif bound to **(A)** the pseudokinase domain of TRIB1, and **(B)** the WD40 domain of COP1. **(C)** Fluorescence polarisation displacement assay of FITC-TRIB1(349–367) from the WD40 domain of COP1 by either TRIB1(84–372) alone at varying concentrations, or TRIB1(84–372) in the presence of 10 μM C/EBPα degron peptide. Points represent the mean of three technical replicates (generated using two separately prepared stocks of COP1), and error bars represent the SEM. **(D)** Summary of molecular dynamics simulations of SLE-out TRIB1, or SLE-in TRIB1 with and without substrate. The number of hydrogen bonds between the C-terminal tail and β4 strand are plotted over the time course of the simulation above, and an indicative state during the latter part of the simulation is shown below. Animated trajectories are available in movie S2.

To further investigate the structural basis of TRIB1 allostery we performed molecular dynamics simulations comparing the dynamics of TRIB1 in its autoinhibited form, in its active conformation but without substrate, and in its active conformation in the presence of substrate. These analyses could be used to: i) visualize the effect of substrate binding on stability of the C-terminal tail of TRIB1, and ii) infer key residues involved in the allosteric regulation of TRIB1.

Firstly, to understand the propensity of the TRIB1 C-terminal tail to occupy its autoinhibited conformation, we analysed the stability of the tail over the parallel simulations (movie S2). In the structure of autoinhibited TRIB1 (PDB: 5cem), the tail stably resides in its binding cleft atop the αC-helix, maintaining multiple hydrogen bonds with the β4 strand. In contrast, initiating a simulation of TRIB1 (SLE-in) bound to C/EBPα with the C-terminal tail in an equivalent position alongside the β4 strand shows the tail rapidly dissociating from its binding site. Beginning a simulation with TRIB1 in its active (SLE-in) conformation but without substrate present, causes the tail to remain bound for an intermediate length of time, but eventually dissociate. To quantify these observations, we plotted the number of hydrogen bonds between the TRIB1 C-terminal tail and the β4 strand over the course of the three simulations (Fig. 4D). This analysis highlights that the C-terminal tail is less effectively sequestered when TRIB1 occupies an active state, and becomes fully destabilised when substrate binding completes the active complex.

Secondly, to understand the key residues inherent to the allosteric mechanism we compared the difference in torsion angle dynamics of each residue in TRIB1 using a Kullback-Leibler (KL) divergence based statistics (See methods; Fig. 5). Quantification of KL divergence of TRIB1 without the C-terminal tail (Fig. 5A) indicates that structural regions (such as the catalytic loop and activation loop) in direct contact with C/EBPα show significant difference between substrate-bound and unbound simulations. Interestingly, the KL divergence profile of TRIB1 with the C-terminal tail included reveals differences in torsion angle dynamics of residues not only in the substrate binding site, but distal regions of the pseudokinase domain, including the C-terminal tail, SLE-motif, G-loop, and αC-helix (Fig. 5A). In the presence of substrate, the SLE+1 aspartate (Asp228) displays large conformational flexibility in that it can make charge interactions with Lys220 from the catalytic loop, Arg102 from the glycine-rich loop and Lys130 from the αC-helix during the simulation (Fig. 5B). Therefore, the SLE-motif seems to be the global hub in propagating the allosteric signals of C/EBPα substrate binding. Together, these data reinforce the key functional role of sequence features that are unique to Tribbles proteins, which contributing to a mode of regulation reminiscent of active kinases.

**Fig. 5.**
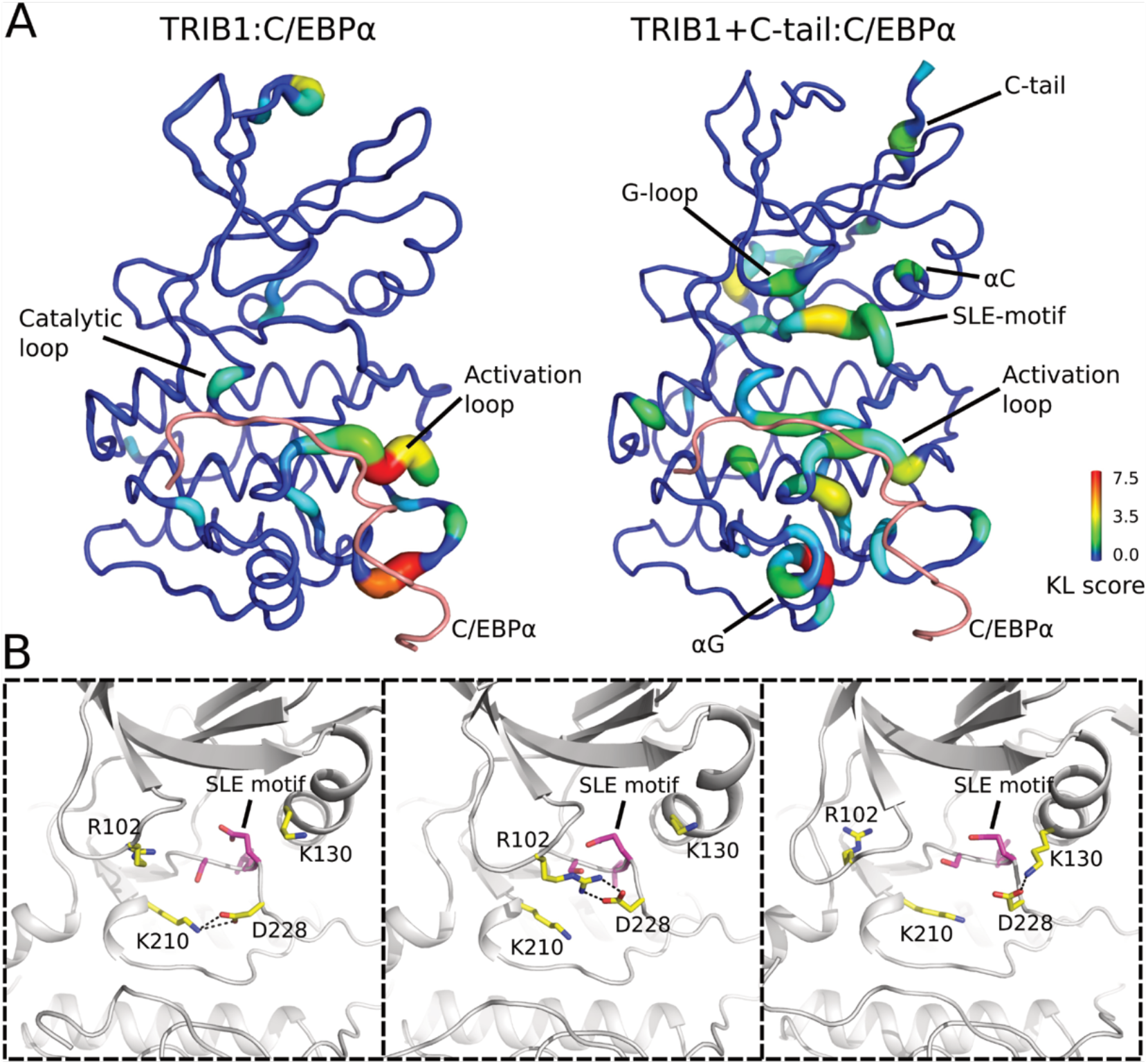
Comparative analysis of molecular dynamics simulations of Trib1 pseudokinase domain with and without C/EBPα peptide. **(A)** The Kullback-Leibler (KL) divergence of the torsion angle of each residue is shown. Motifs and residues that display differential distribution between the C/EBPα unbound simulation and C/EBPα bound simulation are labeled. Left panel: KL divergence of Trib1 simulations without the C-terminal tail. Right panel: KL divergence of Trib1 simulations with the C-terminal tail. **(B)** Representative snapshots from TRIB1+C-tail:C/EBPα simulation. SLE+1 aspartate (Asp228) mediates charge interaction with Lys210 from catalytic loop, Arg102 from glycine-rich loop and Lys130 from αC-helix.

### Potential for nucleotide or small-molecule binding by TRIB1

A clear consequence of conformational changes induced by C/EBPα-binding is the opening of the TRIB1 active site, which in the SLE-out state is blocked by the activation loop (Fig. 6A). In contrast, the SLE-in state induces a large open binding pocket (Fig. 6B). Such an open state would be consistent with reports that the closely-related pseudokinase TRIB2 can bind ATP, and to small molecule ligands (31, 32). We initially tested the effect of C/EBPα degron on the melting temperature of TRIB1(84–372) in a differential scanning fluorimetry (DSF) assay (33). The addition of C/EBPα degron peptide induced a significant stabilisation of TRIB1 (Fig. 6C; comparing orange to grey). However, further addition of 200 μM ATP did not shift the melting temperature of either TRIB1 alone (Fig. 6C; green) or the TRIB1-C/EBPα mixture (Fig. 6C; black) relative to their nucleotide-free equivalents. Thus, it appears that movement of the SLE-motif to its SLE-in conformation is insufficient to confer the ability to bind ATP.

**Fig. 6.**
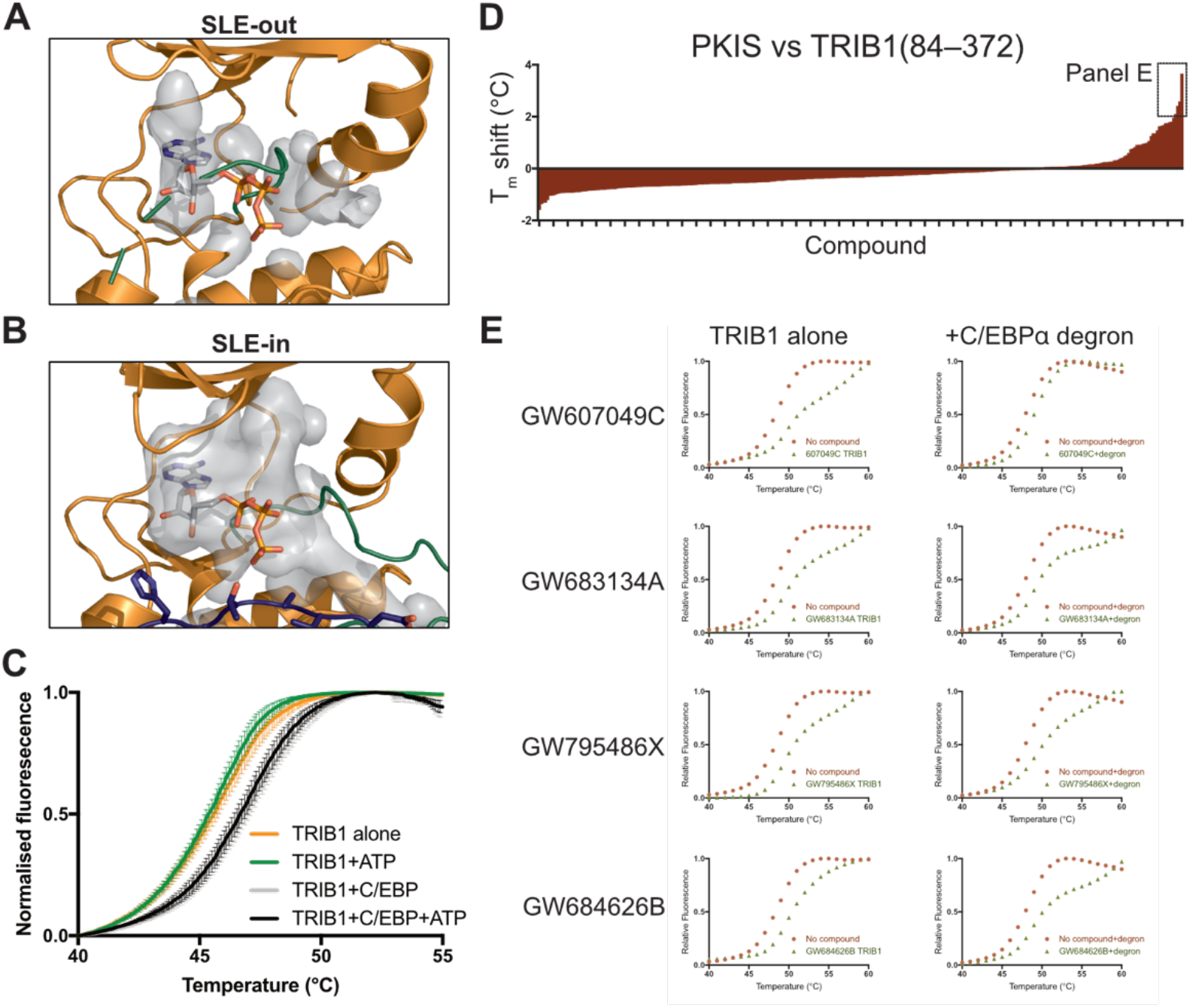
TRIB1 can potentially bind small molecule ligands but not ATP. Representation of the available binding cavity of TRIB1 (grey) in **(A)** the SLE-out conformation from PDB 5cem, and **(B)** the SLE-in conformation when bound to C/EBPα. **(C)** DSF melting analysis of TRIB1 or TRIB1+C/EBPα degron peptide in the presence or absence of ATP. **(D)** Summary of DSF analyses of TRIB1(84–372) against the PKIS library. T_m_ are expressed as difference between each compound and the mean of three DMSO-only controls. The four compounds shown in more detail in panel E are indicated. **(E)** Individual DSF melting curves of four selected compounds (left), with additional titrations for each that include C/EBPα degron peptide (right).

Even though TRIB1 does not appear capable of binding ATP in either conformation, there is no obvious steric hindrance of the rear (adenine-binding) portion of the TRIB1 ATP-binding site, particularly in the SLE-in state. This is relevant because ATP-competitive inhibitors often occupy of the adenine-binding portion of a kinase active site, rather than the phosphate-stabilising regions that are defunct in TRIB1. To explore this possibility we used DSF to screen TRIB1 against the publically available protein kinase inhibitor screen (PKIS; (34)). We found several compounds that induced thermal stabilization of TRIB1 by greater than 2 °C (Fig. 6D). The four hits were originally designed as inhibitors of either TIE2/VEGFR2 or EGFR/ErbB2 and were built upon either the benzimidazolyl diaryl urea, furopyrimidine (both TIE2/VEGFR2-targeting) or anilino thienopyrimidine (EGFR/ErbB2-targeting) scaffold (35–37). Moreover, three of the four also induced similar stabilisation of TRIB1 in the presence of C/EBPα degron peptide whereas GW607049C induced a different melting profile in the presence and absence of degron (Fig. 6E, fig. S7), which could indicate binding to different conformations of TRIB1. Although further characterisation is definitely required, these results offer the first suggestion that inhibitors could be developed to target TRIB1. Based on structural and mechanistic studies presented here, development of specific ligands that bind the SLE-out or SLE-in conformations of TRIB1 could have exciting potential to either block substrate-binding, or release autoinhibition of TRIB1 to promote recruitment of the COP1 ubiquitin ligase.

## Discussion

Tribbles pseudokinases function in concert with the COP1 ubiquitin ligase to degrade critical regulators of metabolism and transcription (4). In this study, we show that TRIB1 recognises conserved degrons in C/EBP family members with similar efficacy, and that binding of C/EBPβ can be antagonised by C/EBPβ phosphorylation. We also report the structure of TRIB1 in complex with the degron from its canonical substrate, C/EBPα. The mechanism of C/EBPα- binding by TRIB1—a pseudosubstrate-like binding mode—couples substrate-binding on one face of TRIB1 to release of the C-terminal tail on the opposing face of the molecule. Such a mechanism seems likely to have important consequences for protecting TRIB1 from unproductive degradation by COP1, but also offers exciting potential when considering ligands designed to bind the TRIB1 active site.

Pseudokinases are not constrained by the need to perform catalysis, and as such have evolved as diverse mediators of signal transduction via three major mechanisms; allosterically activating other enzymes, acting as signalling switches, or scaffolding protein–protein complexes (2, 3). There are few examples of pseudokinase structures captured in both inhibited and ‘active’ states, but the mechanism described here for TRIB1 bears remarkable resemblance to regulation of conventional kinases. However, it is facilitated by unique structural features of TRIB1. For instance, the αC-helix is commonly mobile in conventional kinases (1), but in TRIB1 it seems to be particularly malleable due several proline residues creating a non-ideal helix. Moreover, in addition to possessing an SLE-rather than DFG-motif, TRIB1–3 all possess a glutamate preceding the SLE, which is unique, stabilises the autoinhibited state of TRIB1 and exhibits increased Kullback-Leibler divergence in molecular dynamics simulations of substrate-free and - bound TRIB1-C/EBPα (Fig. 5). Overall, the mechanism described in this work offers an intriguing example of sequence features that are divergent between pseudokinases and kinases maintaining orthologous function as an allosteric switch. In conventional kinases binding of regulators to the αC-helix and stabilisation of the regulatory spine generally facilitates substrate binding and activity (1); in the case of TRIB1 it appears that the reverse is true. Substrate binding promotes conformational changes in the αC-helix that disrupt sequestration of the TRIB1 C-terminal tail, and subsequent complex formation with COP1. Whether similar mechanisms control protein-protein interactions in other pseudokinases, or active kinases, is an interesting proposition.

It has recently been shown that TRIB2 also binds its own C-terminal tail (32), and thus potentially also adopts an autoinhibited conformation that facilitates allosteric regulation. Although no structures are available for TRIB2 or TRIB3, the structure of the STK40 pseudokinase domain was recently reported (30), which adopts a conformation similar to the substrate-bound state of TRIB1 (fig. S6). However, STK40 is the most divergent Tribbles family member and does not share the SLE-type motif or preceding glutamate residue. In this sense it appears unlikely to share the same allosteric switch mechanism as TRIB1, and may be constitutively-active with regards to substrate binding. The impact of such for STK40 stability, and biological functions independent of canonical Tribbles family-members will be an area of some interest.

C/EBP transcription factors play pleiotropic roles in development of adipocytes and myeloid cells. The importance of C/EBPs in myeloid development is illustrated by mutations in C/EBPα that occur in ~10% of acute myeloid leukaemia patients (38), and the critical transcriptional role of C/EBPβ in multiple myeloma (39). Many of the biologically characterised roles of TRIB1 and TRIB2—for instance regulation of lipid metabolism or macrophage development—can be traced to their ability to control levels of C/EBPα in particular. The identification of the Tribbles degron present in the p42, but not the dominant negative p30 isoform of C/EBPα, suggested one mechanism by which Tribbles may selectively control a single transcription factor (17). We now show that the Tribbles degron is conserved in four of six C/EBP family members, of which all but C/EBPδ are similar to C/EBPα in that they also have short isoforms that lack a Tribbles degron. Because Leucine zipper transcription factors can form either homo- or hetero-dimers, which may contain no, one or two Tribbles degrons there appears to be ample opportunity for gradation of C/EBP ubiquitination by COP1-TRIB1/2, where complexes containing two degrons are preferentially degraded by Tribbles, over complexes that might be recruited less avidly. Furthermore, C/EBPβ can apparently be protected from degradation by phosphorylation (Fig. 2D). Such protection may have clinical relevance when considering tumours with constitutively-activated kinases upstream of EBPβ. Namely, given the role of EBPβ in Ras-induced transformation and myeloid cancers it is tempting to speculate that pharmacologically preventing EBPβ phosphorylation could make it susceptible to Tribbles-mediated degradation, and offer benefit in specific cancers (28, 29, 40). Phosphorylation of EBPβ further adds to the complexity of COP1-based ubiquitination and phosphorylation, given that ETS-family transcription factor degradation is reduced upon phosphorylation by Src-family kinases (41, 42).

Although the capability of TRIB2 to bind ATP appears not to be realized in either the SLE-in or SLE-out forms of TRIB1 ((31); Fig. 6), this work opens the intriguing possibility that small molecule inhibitors targeted towards the active site of Tribbles could be used to block function. An exciting recent report suggests that small molecule inhibitors that covalently target a cysteine-rich segment in the αC-helix of TRIB2 can induce its degradation (32). The fact that these inhibitors were also originally designed against EGFR offers some overlap with the binding profile shown here for TRIB1. The chances of such overlap were likely increased by our having screened an identical library to Foulkes et al., which is enriched in EGFR targeted ligands. A more extensive search of chemical space is warranted to find more potent binders of TRIB1. The mechanism revealed in this work offers an exciting incentive for finding Tribbles-targeting small molecules—despite having poor ATP binding propensity the active-site pocket is intimately linked to regulation of TRIB1, and binding of inhibitors around the SLE-sequence and αC-helix may have drastic implications for the fate of TRIB1 or TRIB2 in cells.

Targeting pseudokinases with small molecules is an exciting area of development, for instance: small molecules that stabilize the KSR pseudokinase anatagonize RAF heterodimerization and subsequently Ras signalling (43); and molecules targeting the HER3 pseudokinase modulate heterodimerization with kinase-active EGF-family receptors and induce degradation of HER3 (44). The distinct sequence features within the active site of TRIB1 may offer several possibilities for developing specific inhibitors. It remains a tantalising prospect that high affinity small molecules could manipulate the allosteric mechanism described here to protect substrates, or promote Tribbles degradation, in cancers promoted by Tribbles pseudokinases.

## Materials and Methods

### Protein expression and Purification

All bacterial constructs (TRIB1 and C/EBP) were expressed in *E. coli* BL21 (DE3) cells. COP1 was expressed in insect cells.

#### TRIB1/2 and C/EBP proteins

TRIB1(84–372), TRIB2(53–343) and TRIB1(84–343) linked by a-GSGSSGGPG- linker to C/EBPα(53–75) were cloned into modified pET-LIC vectors incorporating an N-terminal 6 X His tag and a 3C protease cleavage site. C/EBPα(53–75), C/EBPβ(64–86), C/EBPδ(50–72) and C/EBPε(31–53) all fused to maltose-binding protein (MBP) were cloned into modified pET-LIC vectors incorporating an N-terminal 6 X His tag followed by MBP and a 3C protease cleavage site.

Cell pellets were resuspended in purification buffer (50 mM Tris pH 8.0, 300 mM NaCl, 10% (*v/v*) glycerol and 10% (*w/v*) sucrose) supplemented with 0.2 mg/mL lysozyme and 80 U/mL benzonase. Cells were lysed by sonication (Sonifier, Heat Systems Ultrasonics). Protein was initially purified by Ni^2+^ affinity chromatography (His Select, Sigma). Proteins were eluted with purification buffer containing 300 mM imidazole. Protein containing fractions were pooled and digested overnight with 3C protease and 2 mM DTT. The protein was further purified by size-exclusion chromatography using a Superdex 200 Increase column (GE Life Science) (10 mM Hepes pH 7.6, 300 mM NaCl, 0.5mM TCEP) or anion exchange (Resource Q). The core peak fractions were pooled and snap frozen for storage at −80°C.

#### GST pulldowns

Mutations were generated in Trib1(84–372) in a modified pGEX-LIC vector incorporating an N-terminal GST tag using Quikchange mutagenesis. Cell pellets from expression of wild-type and mutant TRIB1 were resuspended in purification buffer and supplemented with 0.2 mg/mL lysozyme and 80 U/mL benzonase and cells were lysed by sonication. The soluble fraction was bound to GST sepharose resin (Amersham Biosciences), and analysed for purity by SDS-PAGE. GST-TRIB1 was incubated with 500 μM C/EBPα(53–75) fused to MBP at 4°C for 20 min using PBS pH 7.4. After incubation the resins were washed with PBS pH 7.4, 0.02% (v/v) Tween-20. Samples were then resuspended in SDS sample buffer and visualised on 12-18% SDS PAGE stained with coomassie R250.

#### COP1 WD40 domain

COP1 WD40 domain (376–731) was expressed in the *Trichoplusia ni* (*Tni*) and *Spodoptera frugiperda* (*Sf9*) cell lines using baculovirus produced in *Sf9* cells. All cells were purchased from Expression Systems, and cultured in ESF 921 medium at 27°C, shaken at 125 r.p.m. The Bac-to-Bac baculovirus expression system (Invitrogen) was used, as per the manufacturer’s instructions, to produce COP1 WD40 baculovirus stocks, except FuGENE 6 (Promega) was used as the transfection reagent. Cells were plated at a density of 8.0 × 10^5^ cells/mL and allowed to grow until logarithmic growth was achieved, as indicated by a density of 1.0 × 10^6^ cells/mL. Subsequently, cells were inoculated with COP1 WD40 baculovirus with an MOI= 1, and incubated at 28°C for 72h (*Sf9*) or 48 h (*Tni*). Cells were harvested by centrifugation at 1000 X g, 5 min. Cell pellets were resuspended in 50 mM Tris-HCl pH 8.0, 10% (*v/v*) glycerol, 150 mM NaCl and 10 mM imidazole supplemented with *DNaseI* (Applichem) and Protease Inhibitor Cocktail (Sigma; p8340). Cells were lysed using an EmulsiFlex-C3, and the soluble fraction purified by Ni^2+^ affinity chromatography (His-Select resin, Sigma). COP1 WD40 protein was eluted from resin in lysis buffer including 500 mM imidazole. The protein was further purified by size exclusion chromatography using a Superdex 200 Increase column (GE Life Science) (150 mM NaCl, 10 mM HEPES buffer pH 7.6).

### Crystallisation and structure determination

TRIB1(84–343) with a GSGSSGGPG linker to C/EBPα(53–75) (~12.2mg/mL) was initially crystallised in 2.8 M Na Acetate (Salt) 0.1 M BIS-TRIS prop 7 pH. Initials hits were refined using additive screening and soaking experiments, and crystals for data collection were grown in 2.8 M Na Acetate 0.1 M BIS-TRIS prop 7 pH with 5% sodium citrate mother liquor diluted to 75 and 80% and drop ratios of 2:1 and 3:1. The crystals were cryoprotected in mother liquor supplemented with 25 % glycerol. In attempts to improve diffraction, the crystal that eventually yielded the best resolution included 1 mM ATP in the cryoprotectant, but there was no electron density attributable to nucleotide in the final structure. Data was collected at the MX2 beamline of the Australian Synchrotron (Table 1), and processed using XDS and AIMLESS (45, 46). The structure was solved by molecular replacement in Phaser using the pseudokinase portion of PDB 5CEM as a model (47). Rebuilding was performed in COOT (48), and the structure was refined against diffraction data to a final resolution of 2.8 Å using Refmac (49).

### Molecular Dynamics

#### Structural Modelling

The SLE-out conformation of TRIB1(88–359) was modelled based on the PDB structure 5CEM (17). Missing residues of the αC-β4 loop(140-141), activation loop (229– 237) and part of the C-terminal tail(343–356) were fixed using the automodel class in Modeller (50). The C-terminal tail(341–364) was incorporated into the SLE-in-TRIB1:C/EPBα complex using PDB 5CEM as the template. The simulations were performed in the presence and absence of C/EPBα peptide.

#### MD System Preparation and Simulation

All MD simulations were performed using Gromacs version 5.1.2 (51). Specifically, the protein was parameterized using amber99sb-ildn force field and solvated with TIP3P water model (52). The protein was centered in a dodecahedron box with the distance between the solute and the box larger than 1nm in all directions. 0.1 mM ions (Na+ and Cl-) were added to the system by randomly replacing solvent molecules to neutralize the charge of the system. Steepest descent and conjugate gradient algorithms were employed in conjunction to minimize the potential energy of the system so that the maximum force (Fmax) is less than 100 kJ/mol/nm. LINCS (Linear Constraint Solver) algorithm was used to constraint bonded interactions. Verlet cutoff scheme is used to maintain neighbor-list (53). Long range electrostatic interactions are calculated based on Particle-Mesh Ewald (PME) method (54). Temperature equilibration was done in canonical ensemble (NPT) for 200 ps using V-rescale thermostat (55). Isothermal-isobaric ensemble (NPT) was then maintained through Berendsen barostat at 1.0 bar for 200 ps. The productive simulations were carried out at NPT ensemble for 1 μs. The torsion angle analysis of the trajectory was calculated using the gmx chi module of Gromacs.

#### Kullback-Leibler (KL) Divergence Analysis

Kullback-Leibler divergence is a statistical measure of how two probability distributions differ from each other. In this analysis, we focused on the distribution of the torsion angles (Φ, Ψ, Χ) of each residue in Trib1 during the MD simulations. We used the C/EBPα unbound form as the reference state and C/EBPα bound form as the target state. MutInf package was used to perform the KL calculation (56). Specifically, we split the 100–1000 ns of our MD data into 4 equal length blocks as the bootstrapping set and discretized the torsion angle (Φ, Ψ, Χ) distribution of each residue with a binwidth of 15 degrees. The sum of the KL divergence of all angles of a given residue was reported as the KL score for that residue. The KL score was then mapped to the Trib1 structure and visualized using the putty representation of PyMOL.

### Biophysical measurements

#### Isothermal Titration Calorimetry (ITC)

ITC experiments were performed at 30 °C using a VP-ITC calorimeter (GE Healthcare). TRIB1(84–372), and MBP fusion proteins of C/EBPα(53–75), C/EBPβ(64–86), C/EBPδ(50–72) and C/EBPε(31–53) were initially purified by Ni^2+^ affinity chromatography followed by size-exclusion chromatography using a matched buffer consisting of 10 mM HEPES (pH 7.6), 300 mM NaCl, 0.5 mM TCEP. C/EBP (200 mM) was injected into 20 mM TRIB1(84–372). Baseline corrections and integration were performed using NITPIC (57), isotherms were fit to a single site-binding model using SEDPHAT (58), and figures were generated using GUSSI (http://biophysics.swmed.edu/MBR/software.html).

#### Fluorescence Polarisation

fluorescence polarisation of synthetic TRIB1 and C/EBPα peptides bearing an N-terminal fluorescein isothiocyanate (FITC) fluorophore (Mimotopes) was measured as indicated in combination with purified COP1 WD40 domain (376–731), TRIB1(84–372), TRIB2(53–343) or C/EBPβ degron peptide. COP1-TRIB1 and TRIB1/2-C/EBPα K_D_ determination were performed in black 384-well microplates (Greiner Bio-One) format with a final reaction volume of 30 μL, while TRIB1-C/EBPα displacement by C/EBPβ was performed in 96-well format with a final reaction volume of 60 μL. For COP1 displacement measurements, the COP1 WD40 domain and TRIB1-FITC peptide (349–367) were kept at a constant concentration of 1.5 μM and 25 nM respectively, diluted in a fluorescence polarisation buffer (10 mM HEPES, 300 mM NaCl, 0.5 mM TCEP and 0.02% (v/v) Tween-20). The concentrations of competing ligand, either TRIB1 protein or TRIB1 peptide were varied from 0–20 μM. Where necessary the reactions were supplemented with C/EBPα peptide (53–75) at a concentration of 10 μM. Once all reactions were prepared they were allowed to incubate at room temperature for 20 minutes and measured using the PolarStar (BMG Tech) plate reader. Following this method, the data from three independent experiments were obtained and plotted as the mean ± S.E.M using Prism 7. For TRIB1-C/EBPα displacement by C/EBPβ, 1.5 μM TRIB1, 25 nM C/EBPα- FITC peptide were incubated together and C/EBPβ was titrated in at concentrations from 43 nM to 350 μM and measurement was performed in an equivalent manner.

#### Differential Scanning fluorimetry

The PKIS screening library in 384-well format was received from SGC-UNC at 10 mM stock concentration. The stock was prediluted to 2 mM, before aliquoting into black 384 well plates (Greiner Bio-One) using the mosquito^®^ LCP (TTP labtech) and screening against TRIB1 at a final compound concentration of 40 μM. TRIB1(84–372) was diluted for use at 5 μM (with or without 25 μM C/EBPα peptide) with DSF buffer (10 mM Hepes pH 7.6, 300 mM NaCl, 0.5 mM TCEP). The protein was incubated in the plate at room temperature for 30 minutes. Sypro^®^ Orange was then diluted with DSF buffer for use at 5X and was pipetted into each well. The plate was then covered with PCR film and centrifuged for 5 minutes before being measured on a Roche lightcycler^®^ 480 using the default Sypro^®^ Orange protein programme. The data was initially condensed using R version 3.4.3. Data was then analysed in Microsoft Excel using the CS example DSF Analysis v3.0 template provided by the Structural Genomics Consortium (59, 60), and Boltzmann fitting in Graphpad Prism 7.

## Acknowledgments

This research was undertaken in part using the MX2 beamline at the Australian Synchrotron, part of ANSTO. We thank the New Zealand synchrotron group for facilitating access to the MX beamlines. This study was supported in part by resources and technical expertise from the Georgia Advanced Computing Resource Center, a partnership between the University of Georgia’s Office of the Vice President for Research and Office of the Vice President for Information Technology. We thank Patrick Eyers (University of Liverpool) for provision of template to create TRIB2 expression constructs.

## Data and materials availability

The structure of the TRIB1-C/EBPα is deposited in the protein databank with accession code XXXX.

---

**Supplementary Figure 1.**
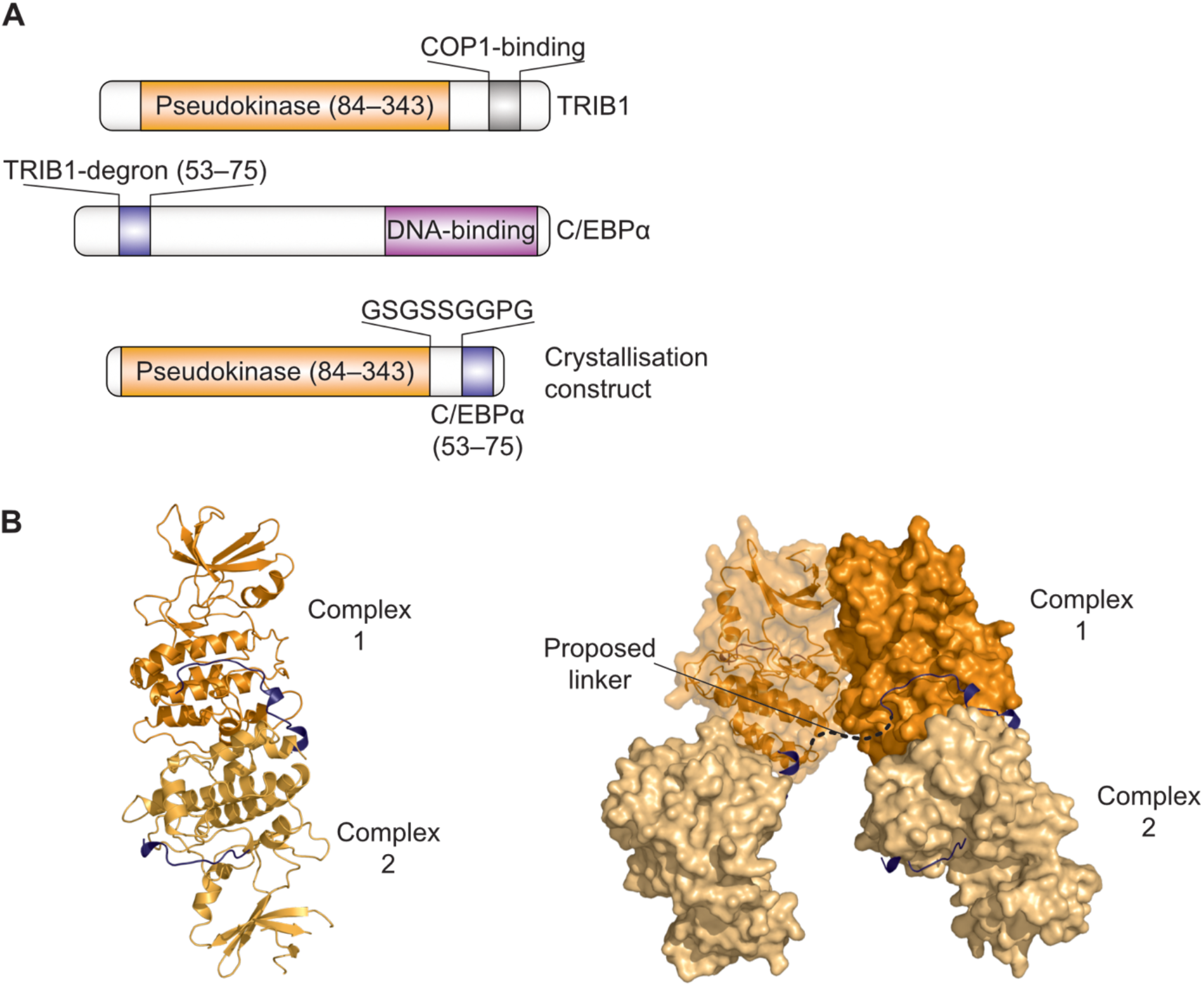
Construct Design for crystallization. (A) Schematic illustration of the fusion construct that allowed successful crystallisation of the TRIB1-C/EBPα degron complex. (B) Illustration of the crystal packing within the unit cell. The linker between the TRIB1 pseudokinase and C/EBPα is not defined (dotted line), and could arise by C/EBPα being contributed by a TRIB1 molecule across a side-by-side packing or a packing between C-terminal lobes.

**Supplementary Figure 2.**
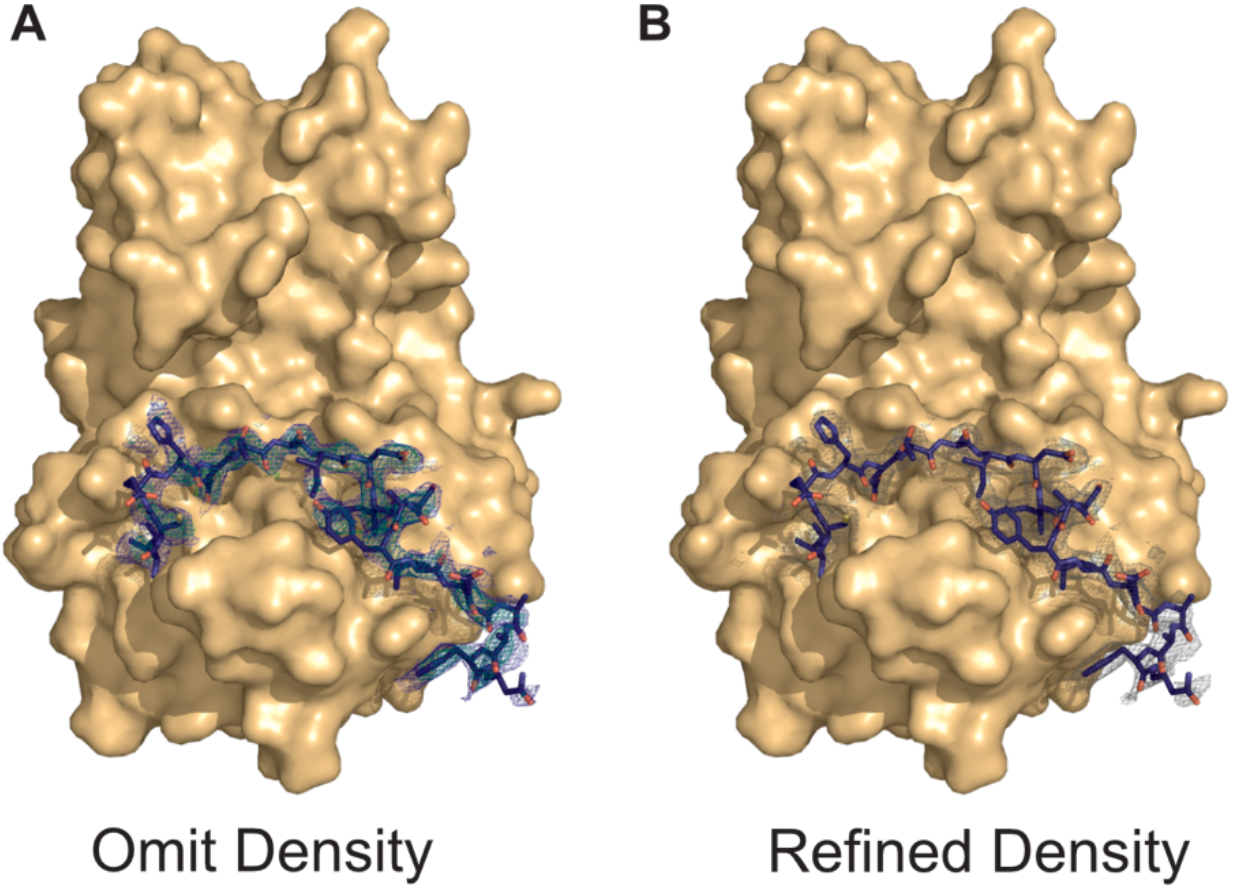
C/EBPα degron electron density. (A) Electron density maps (2Fo-Fc blue, Fo-Fc (green)) contoured to 1.0 and 3.0 σ respectively, following a cycle of model refinement omitting the C/EBPα peptide. (B) 2Fo-Fc electron density map contoured to (1.0 σ) after refinement of the final structure.

**Supplementary Figure 3.**
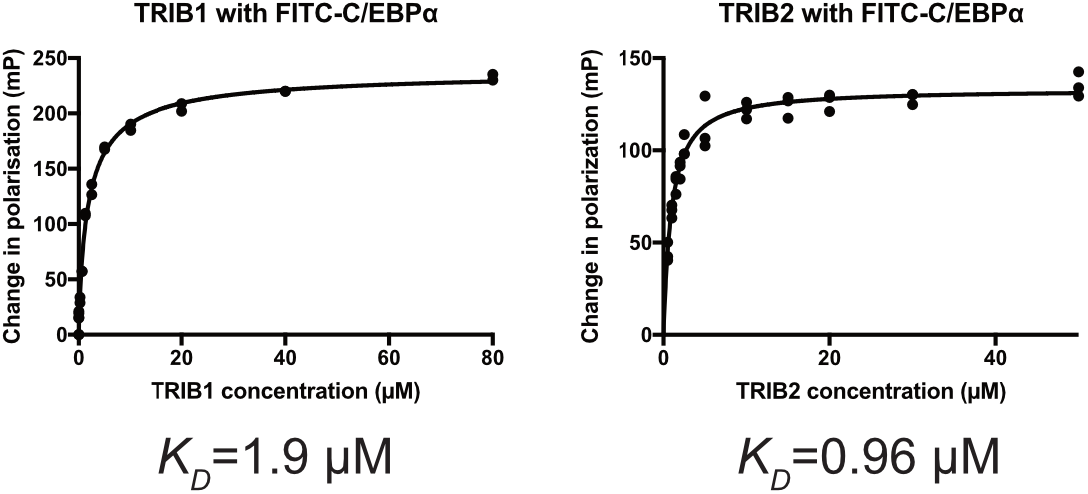
Dissociation constant of TRIB1/2 binding C/EBPα. Measurement of the dissociation constant of the TRIB1-C/EBPα degron monitored using fluorescence polarisation of FITC-C/EBPα degron peptide (25 nM) and varying concentrations of TRIB1(84–372) or TRIB2(53–343).

**Supplementary Figure 4.**
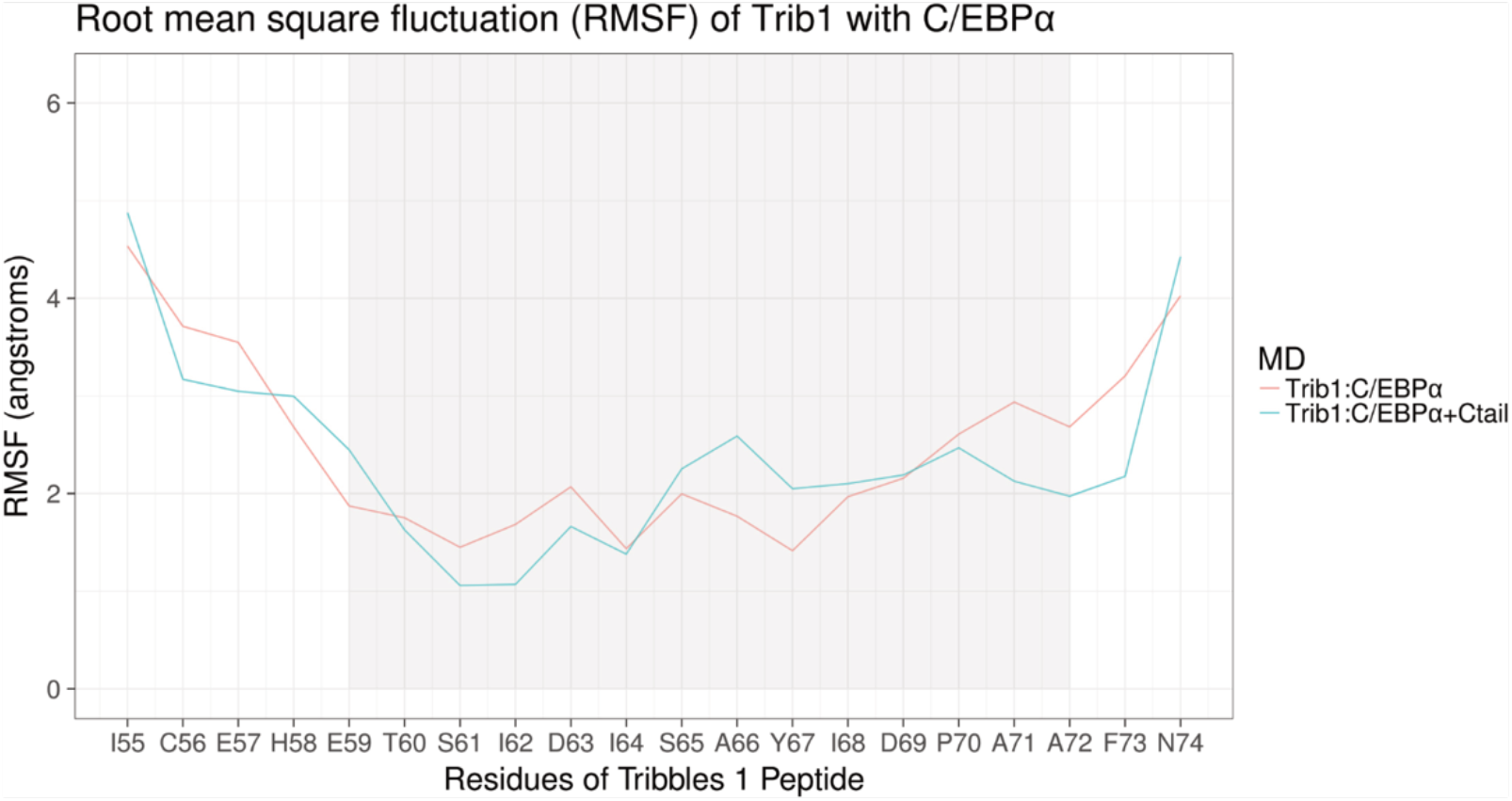
Stable regions of C/EBPα degron in complex with TRIB1. RMSF plot of C/EBPα residues during 1 μs simulation in complex with TRIB1. Simulations were performed both incorporating (red) and omitting (blue) the C-terminal tail of TRIB1 (see also Fig. 4D), indicating that the C-terminal tail of TRIB1 does not impart a significant stabilising or destabilising effect on substrate binding.

**Supplementary Figure 5.**
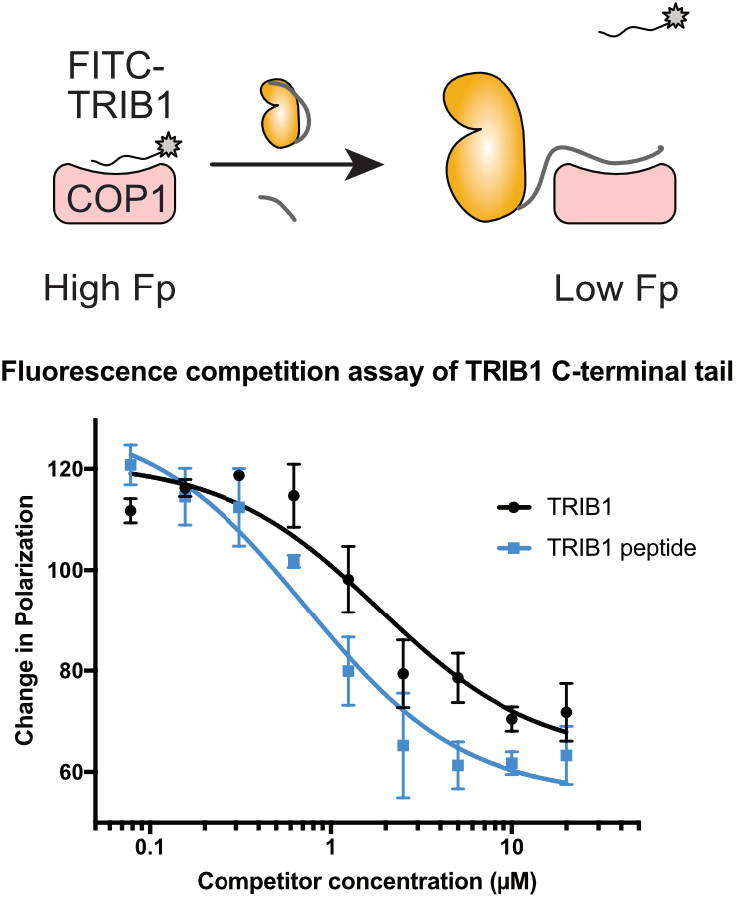
Displacement of the TRIB1 C-terminal tail from COP1. Fluorescence polarisation displacement assay of FITC-TRIB1(349–367) from COP1 WD40 domain, by unlabelled TRIB1(349–367) (blue) compared to TRIB1(84 to 372) (black).

**Supplementary Figure 6.**
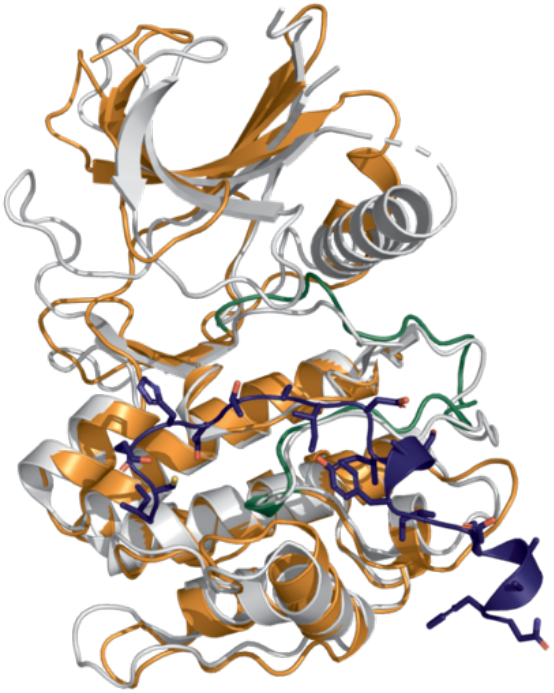
Comparison of TRIB1 to STK40. Overlay of substrate-bound TRIB1 (Orange) with STK40 (light grey; PDB 5L2Q

**Supplementary Figure 7.**
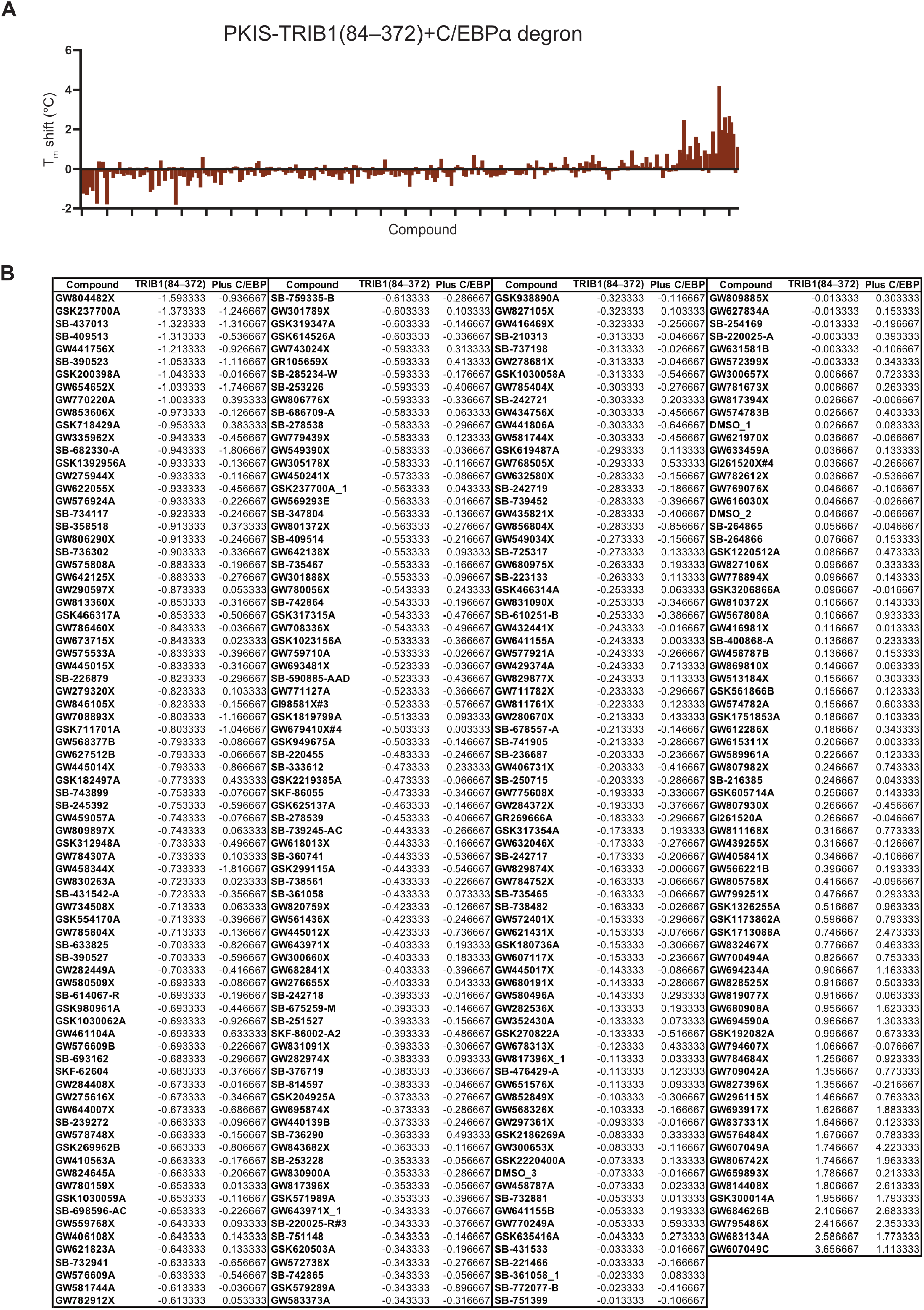
PKIS screening of TRIB1. Differential scanning fluorimetry (DSF) screening of TRIB1(84 to 372) against the PKIS library. (A) Tm shift data, expressed relative to DMSO-only control, for TRIB1-C/EBPα degron peptide mixture. Samples are shown in the same order as Fig. 6D, and generally exhibit a similar binding profile with and without C/EBPα, with some exceptions. (B) DSF data for individual compounds, relating to Fig. 6D and fig. S7A.

Movie S1. Morph between substrate-free (SLE-out) and C/EBPα-bound (SLE-in) structures of TRIB1. Morph was created in PyMOL.

Movie S2. Simulations of the TRIB1-C-terminal tail. Pseudokinase-C-terminal tail interaction during molecular dynamics simulations of TRIB1 SLE-out (left), SLE-in without C/EBPα (centre), and SLE-in with C/EBPα (right).

